# Nanoscale structural response of biomimetic cell membranes to controlled dehydration

**DOI:** 10.1101/2023.06.26.546525

**Authors:** Emilia Krok, Henri G. Franquelim, Madhurima Chattopadhyay, Hanna Orlikowska-Rzeznik, Petra Schwille, Lukasz Piatkowski

## Abstract

Although cell membranes in physiological conditions exist in excess of water, there is a number of biochemical processes, such as adsorption of biomacromolecules or membrane fusion events, that require partial or even complete, transient dehydration of lipid membranes. Even though the dehydration process is crucial for understanding all fusion events, still little is known about the structural adaptation of the lipid membranes when their interfacial hydration layer is perturbed. Here, we introduce the study on the nanoscale structural reorganization of the phase-separated, supported lipid bilayers (SLBs) under a wide range of hydration conditions. Model lipid membranes were characterized with the combination of fluorescence microscopy and atomic force microscopy, and crucially, without applying any chemical or physical modifications, that so far have been considered to be indispensable for maintaining the membrane integrity upon dehydration. We revealed that decreasing hydration state of the membrane leads to an enhanced mixing of lipids characteristic for the liquid-disordered (L_d_) phase with those forming liquid-ordered (L_o_) phase. This is associated with a 2-fold decrease in the hydrophobic mismatch between the L_d_ and L_o_ lipid phases and a 3-fold decrease of line tension for the completely desiccated membrane. Importantly, the observed changes in the hydrophobic mismatch, line tension, and miscibility of lipids are fully reversible upon subsequent rehydration of the membrane. These findings provide deeper insights into the fundamental processes such as cell-cell fusion that require partial dehydration at the interface of two membranes.

## 1 Introduction

Biological cell membranes are permeable barriers, responsible for maintaining homeostasis and protecting the cell from the surrounding environment^1^. They act as gateways, mediating the selective transportation of ions, and biomacromolecules such as glucose or amino acids^2^. Finally, they play a key role in cell compartmentalization, allowing mutually exclusive biochemical processes to occur at the same time within the cell^3^. The study of cellular membranes in their native form is very challenging due to the high structural complexity of these systems as well as the abundance of chemical, biological and physical processes occurring within the cell. For this reason, model biological cell membranes are often deployed, such as giant unilamellar vesicles (GUVs) or supported lipid bilayers (SLBs), which possess analogous physical and structural properties as native cell membranes but at the same time can be modified and simplified to focus on specific biophysical features^4–9^.

The lateral organization of lipid membranes is driven by the interplay of lipid-lipid and lipid-protein interactions^10^. However, this biological system is not complete without the water, presence of which is considered to be an indispensable factor that modulates the structural organization of membranes, phase separation, as well as spatial arrangement of transmembrane proteins, both in model membrane systems and in living cells^11–13^. The water acts on the membrane particularly in the presence of the so-called hydrophobic mismatch, which takes place when the thickness of different membrane constituents is different, leading to an exposure of the hydrophobic moieties to water^14^. This hydrophobic interaction is energetically unfavorable and as a consequence it is one of the main mechanisms driving the phase separation^15^. Garc*í*a-Sáez et. al. modified the thickness of the disordered phase using phosphatidylcholines of different acyl chain lengths and showed that the bigger the difference between the hydrophobic parts of the membrane, the stronger is the phase separation, and the higher is the line tension at the boundary of the two phases^16^. Furthermore, line tension is responsible for the self-healing properties of the cell membranes, facilitating the closure of transient pores within the structure of lipid bilayer^17^. Line tension for single component membranes can be determined by inducing formation of transient pores in the membrane structure and observation of the pores closing rate.

Srividya et al. showed that line tension in single phase membranes increases with the acyl chains length of the constituting lipids^18^.

Although lipid membranes in natural conditions indeed exist in excess of water, there are biological processes that require at least partial, interfacial dehydration of lipid bilayers. Processes such as endo- and exocytosis^19^, neurotransmission^20^, viral entry^21^, fertilization^22^, or cell fusion during embryogenesis^23^ and morphogenesis^24^ involve merging of two lipid membranes. All of these membrane fusion events rely on interactions between lipids, proteins, and water hydrating the two interfacing membranes^25^. When the two bilayers come in close contact and the distance between them is reduced to approximately 2-3 nm, a strong hydration repulsion, also known as “hydration force“ occurs between their hydrophilic surfaces^26–28^. At this stage the thin water layer separating the membranes needs to be expelled in order for the hemi-fusion to take place.

Thus, to fully understand the intricate interactions between membrane constituents and the membrane hydration layer, as well as the biophysical consequences of these interactions, it is crucial to gain insights into the structural properties of cellular membranes when their hydration state is altered. Indeed, Chiantia et al. reported on the structure of SLBs composed of DOPC/SM/cholesterol in complete dehydration conditions using atomic force microscopy (AFM)^29^. They showed that without the presence of stabilizing agents such as trehalose, upon abrupt dehydration and subsequent rehydration, membranes lose their integrity and severe structural damage in the form of holes, aggregates and delimitation of membrane is observed^30^. Similar results were reported by Iriarte-Alonso et al. for single component DOPC membrane, where abrupt dehydration led to the formation of multiple defects and holes and consequently loss of membrane continuity^31^. Many attempts have been made to maintain the structure of desiccated lipid membranes and prevent them from rapid vesiculation. Among them we can distinguish the use of saccharides that are known for increasing the spacing between lipids in dry conditions, thus preventing them from collapsing^30,32^, modification of lipid headgroups to improve the interactions between the lipids and the solid support^33–35^ or introduction of physical confinement, which prevents the interfacial peeling force from destructive membrane delamination^36^. All of these approaches, although successful in membrane preservation, involve alteration of native properties of lipid membranes due to the introduced chemical or physical modifications of either membrane or the solid support. Recently, we presented a novel method of membrane preservation under dehydration conditions, based on the controlled, steady decrease of environmental humidity, which opened up new possibilities to study the behavior and properties of membranes without altering their chemical composition or physical features^37,38^.

In this study, using a combination of fluorescence and atomic force microscopy (AFM), we studied the nanoscale structural response of the phase-separated SLBs to a wide range of hydration conditions. Our results prove that when the dehydration process is carried out in a gradual, controlled manner, the structure of lipid membranes can be preserved even under complete desiccation conditions, without the use of stabilizing agents. The applied here dehydration method allowed maintaining of the overall structural organization of the membrane, without the occurrence of defects or holes. At the same time the removal of bulk water led to a prominent, nanoscale structural reorganization within the membrane. We observed that the process of dehydration causes a significant decrease in the hydrophobic mismatch between the L_d_ and L_o_ phases and as a consequence lowers the line tension at their boundary. Importantly, this process is fully reversible and upon subsequent rehydration, the height mismatch increases to its initial state. We show that the removal of bulk water leads to the extensive mixing of the liquid-disorded (L_d_) and the liquid-ordered (L_o_) phase lipids, and changes to the borderline of the L_o_ phase domains. Last but not least, the presented study employs pioneering methodology of AFM measurements under controlled humidity, which can be applied for studying other model cell systems in varying hydration conditions.

## 2 Materials and Methods

### 2.1 Materials

1,2-Dimyristoleoyl-sn-glycero-3-phosphocholine (14:1 PC or DMoPC), egg yolk sphingomyelin (SM), and cholesterol were purchased from Avanti Polar Lipids, Alabaster AL., USA. Monosialoganglioside (GM1) from bovine brain, 1,2-Dioleoyl-sn-glycero-3-phosphoethanolamine labeled with Atto 633 (DOPE-Atto 633), sodium hydroxide (NaOH), calcium chloride (CaCl2), and sodium chloride (NaCl) were purchased from Merck KGaA, Darmstadt, Germany. Alexa Fluor 488 conjugated with cholera toxin B subunit (CTxB-Alexa 488) was obtained from Molecular Probes, Life Technologies, Grand Island, NY, USA. N-2-Hydroxyethyl piperazine-N’-2-ethane sulphonic acid (HEPES PUFFERAN) was obtained from Carl Roth GmbH & Co KG, Karlsruhe, Germany. All the materials and reagents were used without further purification. Optical adhesive glue Norland 68 was purchased from Norland Products Inc., Cranbury, NJ, USA. The ultrapure water was obtained by using Milli-Q Reference Water Purification System from Merck KGaA, Darmstadt, Germany.

### 2.2 Vesicles preparation

Multilamellar vesicles (MLVs) were formed by dissolving DMoPC, SM, and cholesterol in chloroform at the molar ratio 1:1:1 with the addition of 0.1 mol% of DOPE-Atto 633 dye, resulting in a 10 mM solution of the lipids. For fluorescence imaging purposes 0.1 mol% of GM1 was added for further labelling with CTxB-Alexa 488. The lipid mixture was dried for 20 min under nitrogen gas leaving a thin film of lipids deposited on the bottom of the vial. The dried lipid mixture was further desiccated in a vacuum-dry chamber for at least 2 h to ensure the complete removal of the organic solvent. The lipids were resuspended in the buffer solution (10 mM HEPES and 150 mM NaCl, pH adjusted with NaOH to 7.4) and exposed to four cycles of heating on the hot plate at 60 °C and vortexing. Each heating and vortexing step was performed for 1 min. 10 μL aliquots of the lipid suspension containing MLVs were distributed into new sterilized glass vials and stored at -20 °C for further use.

### 2.3 SLBs preparation

SLBs were formed using previously reported method^39^. In a nutshell, lipid vesicles were diluted 10 times to the final lipids concentration of 1 mM by adding the HEPES buffer. Aliquots containing MLVs were bath-sonicated using Bransonic 1800 ultrasound bath for 10 min to generate small unilamellar vesicles (SUVs). To prepare the solid support for lipids deposition, a thin layer of freshly cleaved mica was glued by UV-activated glue onto a round glass coverslip. A half-cut 2 ml Eppendorf tube was placed on top of the coverslip and sealed with silicone to form a temporary water reservoir, necessary for SLB formation, incubation and washing. 100 μL of SUVs solution was deposited on top of the mica. 2 μL of 0.1 M CaCl_2_ was added to enhance the bursting of the vesicles, which was followed by the addition of 600 μL of buffer (10 mM HEPES and 150 mM NaCl). The additional labelling of L_o_ phase was performed only for confocal imaging by addition of 9 μL of 0.01 mM CTxB-Alexa 488. The sample was incubated for 40 min and then washed to remove the excess vesicles with a total of 20 ml of Milli-Q water. Unless explicitly stated washing was done by using Milli-Q water instead of HEPES buffer, to avoid potential formation of salt crystals on top of the membrane upon dehydration. After the final washing step, Eppendorf tube was gently removed and the sample was transferred to the AFM holder, which then was filled with Milli-Q water, to enable proper hydration. For the measurements at varying hydration conditions, the bulk water was gently removed using pipette and the sample was mounted in the coverslip holder. The nitrogen gas with >90% RH was immediately purged through the two ports of the AFM holder.

### 2.4 Hydration control

The control over membrane hydration state was done by using home-build control unit as previously described^37^. The relative humidity (RH) of nitrogen gas was adjusted and maintained by mixing wet (saturated with water vapor, 95% RH) and dry (∼5% RH) N_2_ gas. The final relative humidity and temperature of N_2_ gas were continuously monitored using the electronic thermohygrometer with 0-95% RH range and 1% precision. Samples were exposed to RH of 90% (62 × 10^19^ water molecules/min), 70% (48 × 10^19^ water molecules/min), 50% (34 × 10^19^ water molecules/min), 30% (20 × 10^19^ water molecules/min), and to completely dry conditions (∼5% RH) during dehydration and rehydration cycles. Here, it should be noted that while relative humidity of the environment in which SLB is placed is not a direct property of the lipid membrane, throughout the manuscript we present our results in terms of RH because this is the parameter that we directly controlled in the experiments. This being said, environmental RH can be directly translated into a hydration state of the lipid membrane expressed in a number of water molecules per lipid molecules as showed in earlier works^37,40,41^.

### 2.5 AFM imaging

AFM measurements were performed using a NanoWizard III system from JPK Instruments, Berlin, Germany, mounted on Zeiss LSM 510 Meta fluorescence microscope. The measurements were done using Biolever Mini cantilever (BL-AC40TS-C2) from Olympus, Tokyo, Japan. Images were acquired in contact mode, using a silicon tetrahedral tip with a 10 nm radius and spring constant of 0.09 N/m. The scan rate was set to 1-2 Hz. The force was maintained at the lowest possible value. Both topography and deflection (error) signals were measured simultaneously for trace and retrace direction. Images were post-processed by applying line-fitting, which corrects for the offset within the image, using JPK processing software from JPK Instruments, Berlin, Germany. Final image analysis was done using ImageJ^42^ and Gwyddion software^43^. Height mismatch between phases was determined by analyzing height distribution histograms.

### 2.6 Line tension calculation

Line tension was calculated based on the theoretical model developed by Cohen et al., in which the line tension is directly related to the height mismatch between the L_o_ and L_d_ phase^14^:

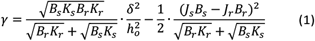

where

γ is the line tension, *δ* is the phase height mismatch, *h*_*o*_ is the monolayer thickness, *B* is the elastic splay modulus, *K* is the tilt modulus and *J* is the spontaneous curvature of the monolayer. The monolayer thickness is defined as an average thickness of L_d_ (*h*_*s*_)and L_o_ phase (*h*_*r*_) monolayers:

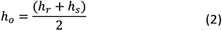

Following the model assumptions presented by Garcia-Sáez et al., we considered that the values of *B*_*r*_ = *B*_*s*_ = 10*kT, k*_*r*_ = *k*_*s*_ = 40*mN*/*m*, and *J*_*r*_ = *J*_*s*_ = 0, that are describing the scenario of “soft“ domains^16^. We determined that the thickness of DMoPC bilayer (L_d_ phase) measured at 90% RH was 3.87 ± 0.21 nm, while the thickness of the L_o_ phase was calculated by adding the value of height mismatch between the two phases to the height of the L_d_ phase.

### 2.7 Confocal imaging

The confocal imaging for sample localization before AFM measurements was done on Zeiss LSM 510 Meta Carl Zeiss, Jena, Germany, using 20x, 0.75NA objective. Confocal images were obtained by using the excitation light of He-Ne laser at 633 nm for Atto 633 and Ar laser was used for excitation of Alexa Fluor 488. Emission was collected in the wavelength range 645-797 nm for the red light channel (Atto 633 detection), and 495-530 nm for the green light channel (Alexa Fluor 488). High-quality images were obtained by using Zeiss 710 microscope with 40x 1.3 NA oil immersion objective. In all imaging experiments minimal laser power was used to minimize photobleaching. To quantify the shape of the lipid domains based on the confocal images obtained at different hydration states the circularity parameter was calculated, defined as:

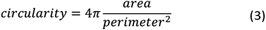

Image processing and calculation of domains circularity were done using ImageJ/Fiji software^44^.

## 3 Results and Discussion

### 3.1 Analysis of fluorescence in dehydrated lipid membranes

Nanoscale characterization of the supported lipid bilayers (SLBs) in different hydration states was done for membranes reconstructed from ternary lipid mixture of DMoPC/SM/cholesterol in the molar ratio 1:1:1. This membrane composition leads to the formation of the L_o_ domains enriched in sphingomyelin and cholesterol, embedded within the L_d_ phase composed of more loosely packed PC lipids. The structure of the membranes with this lipid composition has been well characterized in the literature concerning the size and shape of the domains^16^ as well as membrane dynamics^45^ under full hydration conditions, where the membrane is embedded in an aqueous environment. In contrast, here we focused on elucidating the structural properties of lipid bilayers in a wide range of hydration states: from full hydration (bulk hydration) to dry membrane containing just a few water molecules per lipid. The level of membrane hydration was tuned and controlled using the previously described humidity control setup, schematically shown in Fig. 1A, which purges the water-enriched nitrogen directly to the membrane (see Materials and Methods)^37^. The coupling of confocal microscope with AFM system allowed the simultaneous observation of microscale and nanoscale changes in the lateral organization of the membrane as a function of its hydration state. The analysis of fluorescence images showed that the overall membrane structure remained unaffected by dehydration. Despite the small number of vesicles and aggregates that deposited on top of the bilayer during dehydration, the membrane stayed intact with easily distinguishable phase separation.

**Figure 1.**
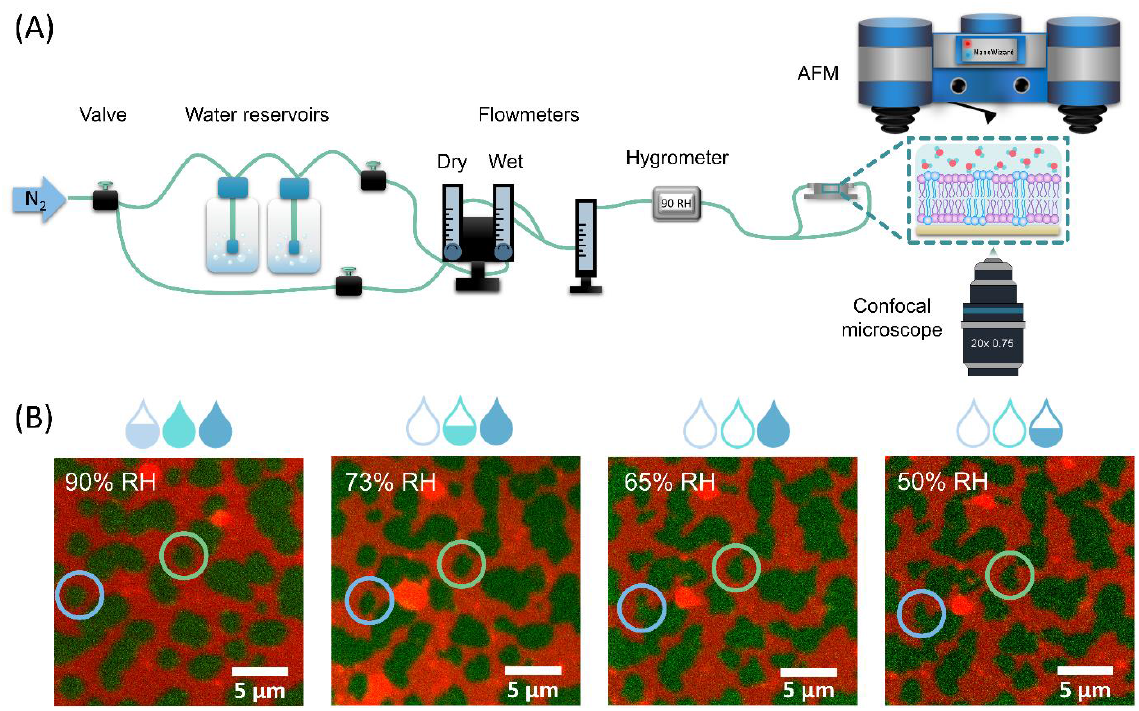
(A) Schematic representation of the home-built humidity-control setup, allowing simultaneous AFM and confocal measurements of biomimetic cell membranes under varying hydration conditions. The set-up consists of a nitrogen gas (N2) source, two reservoirs with water, 3 flow meters, providing information about the dry, wet and final flow reaching the sample, 3 manual valves for precise control of the flow rate, and an electronic thermohygrometer. (B) Fluorescence images of the representative SLB exhibiting phase separation into Ld (labeled with Atto-633-DOPE, shown in red) and L_o_ (labeled with CTxB-Alexa488, shown in green) domains, in different hydration conditions of: 90% RH, 73% RH, 65% RH, and 50% RH. Domains of L_o_ phase become less circular with decreasing hydration level (see for instance domains indicated by blue and green circles). Experiments were performed in 10 mM HEPES and 150 mM NaCl buffer.

However, as the hydration of the membrane is lowered small changes in the shape of L_o_ domains are revealed in a form of more jagged perimeter of the L_o_ phase domains as shown in Fig. 1B. To provide quantitative information about these differences in shape of domains we calculated the circularity parameter, analyzing all clearly resolvable domains from at least four images of two different samples. The domain circularity decreased with the lowering of humidity as shown in Fig. 2A. At fully hydrated conditions domain circularity was 0.82 ± 0.02, while for 5% RH it dropped to 0.51 ± 0.01. It should be noted that the subsequent restoring of high membrane hydration led to the resurgence of domains circularity. The domains circularity values corresponding to the specific hydration levels were the same for the dehydration and rehydration trajectory within the range 5 - 70% RH. However, at 90% RH during the rehydration cycle, the circularity of the domains decreased to the value of 0.53 ± 0.03 (see purple triangle in Fig. 2A). As presented in Fig. S1A and B we observed that at this value of environmental humidity domains were extensively merging, and forming elongated structures composed of few integrated domains, which led to the significant decrease of the circularity parameter. Consistently with our previous reports, at high humidity conditions lipids regain their initial mobility, which promotes merging of the lipid domains37. Apart from the global analysis (of the entire images) of the L_o_ phase circularity at 90% RH, we analysed also 25 domains that did not undergo the merging (see Fig. S1C) and observed that their circularity regained the initial value upon rehydration.

The observed changes in the shape of the L_o_ domains point in the direction of increased miscibility of lipids composing both phases. If indeed true this could also be manifested in the fluorescence signal as one would expect higher L_d_ label fluorescence intensity within the L_o_ phase, due to lipid admixing. To monitor if the compositional changes occurred within domains of the L_o_ phase we measured the average fluorescence intensity of the L_d_ phase labelling dye DOPE-Atto 633 within the domains (I_D_) and divided it over the average intensity of this dye at 1 μm distance from the domain’s border (I_N_). Interestingly, we observed over a 3-fold increase in the ratio of I_D_/I_N_ for the membranes equilibrated at 90, 70, 50, 30, and 0% RH compared to the membrane containing bulk water as shown in Fig. 2B. Upon dehydration, there was a clear increase in fluorescence intensity of DOPE-Atto 633 within domains. The observed increase in fluorescence may result either from the increased migration of lipids from the L_d_ phase into the L_o_ phase or from the changes in photophysical properties of the label fluorophore. It should be pointed out that Atto 633 is characterized by a very high photostability and quantum yield of around 64%. Thus, the observed 3-fold increase of the fluorescence intensity could not result from the changes in the probe quantum yield and rather originated from the diffusion of lipids carrying Atto dye from disordered to ordered phase. The analysis of domains circularity and fluorescence intensity within the L_o_ domains consistently point toward the enhanced lateral reorganization of the membrane constituents under dehydration conditions. The size of the ordered phase domains in this study was around 1-5 μm^2^, thus the details of the structural reorganization are hidden under the resolution limit of fluorescence microscope and could not be resolved based on the fluorescence signal.

**Figure 2.**
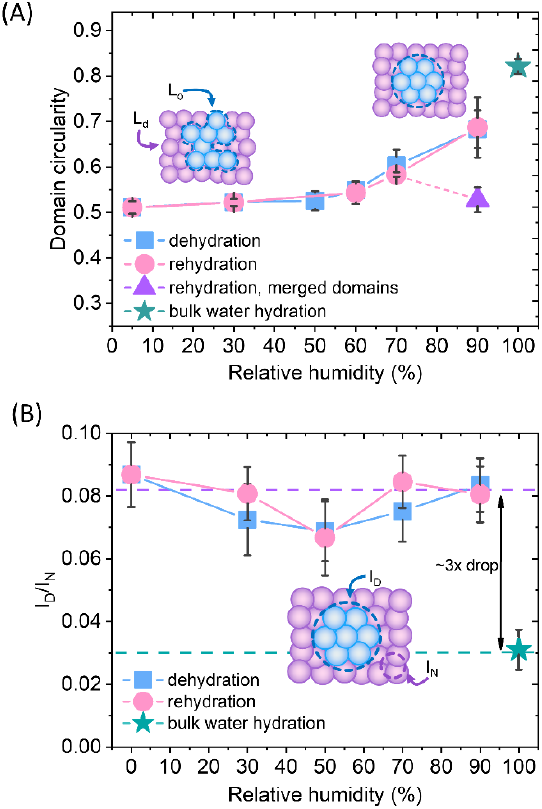
(A) Circularity of the L_o_ phase domains calculated based on the confocal images for fully hydrated sample, during cycle of dehydration (90, 70, 60, 50, 30, 5% RH) and subsequent rehydration (5, 30, 60, 70 and 90% RH). Violet triangle corresponds to the domains circularity calculated for 25 unmerged domains from three different areas. (B) Fluorescence intensity within domain (ID) over the intensity in the area near the domain (IN) as a function of membrane hydration. Experiments were performed in buffer containing 10 mM HEPES, 150 mM NaCl.

### 3.2 Nanoscale structural changes under dehydration conditions

The macroscopic structural analysis of the SLBs in different hydration conditions points toward structural rearrangement of lipids on the nanoscale. To identify the exact origin of the observed changes within the lipid domains, we deployed AFM imaging. The home-built humidity setup was connected directly to the AFM JPK coverslip holder, through perfusion inlets allowing for continuous gas flow inside the chamber as shown in Fig. 1A. In excess of water, we observed well-defined phase separation as shown in Fig. 3A (bulk water). The unlabeled areas measured by confocal microscope corresponded to the more protruding regions detected by AFM and are identified as L_o_ phase lipid domains^46^. Lipid domains in fully hydrated conditions had a round shape with smooth edges. Upon removal of bulk water and subsequent gradual decrease of membrane hydration, the borders between the L_o_ and L_d_ phases became jagged as shown in Fig. S2 (90-5% RH). Simultaneously, we observed formation of small areas of lower height, which we ascribed to the L_d_ phase nanodomains that formed within the L_o_ phase domains, reminiscent of admixing phenomena recently observed for similar mixtures of photoswitchable lipids^47,48^. The measurements right after the removal of bulk water required constant adjustment of the applied force and performing of many scans to remove aggregates deposited on top of the membrane. Moreover, at high humidity membrane is very sticky, it attaches to the AFM tip during scanning, which can drag membrane’s pieces as shown in Fig. S3. Consequently, the measurements at high humidity levels required constant changing of the measured spot, thus during dehydration cycle (Figure S2) we did not follow the structural reorganization of the membrane in the exact same area but rather focused on the global characterization of the sample.

**Figure 3.**
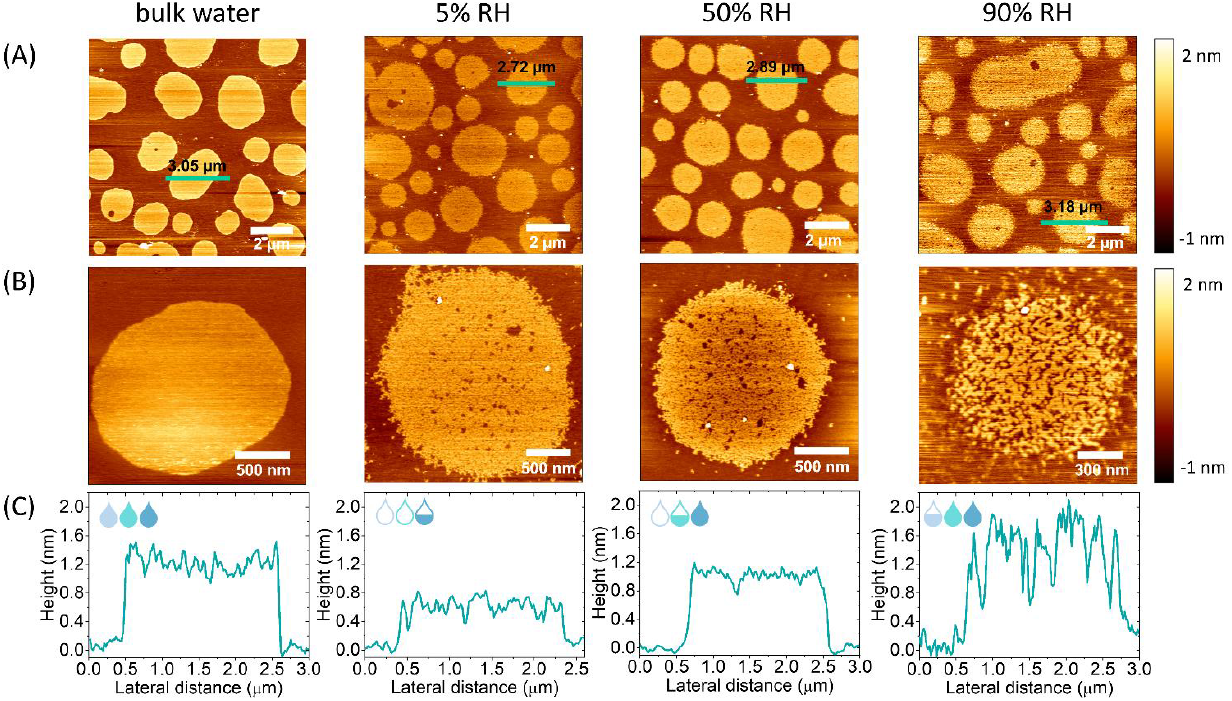
(A) Representative images of fully hydrated SLB, and SLBs after removal of bulk water and equilibrated with 5, 50 and 90% RH. (B) High resolution images of single domains at different hydration levels (bulk water hydration, 5, 50 and 90% RH). (C) Height profiles corresponding to the black horizontal lines shown in (A).

It should be emphasized that the gradual decrease of the humidity did not lead to as prominent reorganization of the domains shape (domain borderline) over the whole cycle of dehydration and rehydration, as it was visible in the case of confocal imaging. From our previous studies, it is clear that upon a decrease of the hydration, diffusion of lipids, as well as their mobile fraction, abruptly drops down when the number of water molecules hydrating the membrane decreases below a certain value^37,49^. At 50% RH and less, the mobility of lipids is almost completely ceased, which results from the breaking of the first hydration shell surrounding a single lipid head group moiety. Furthermore, other recent studies of ours showed that the dynamics of lipids under dehydration conditions is strongly affected by the ionic composition of the buffer hydrating the membrane^50^. Although lipids in the presence of Na^+^ ions remain mobile in humidity above 50% RH, in MilliQ water upon dehydration to 85% RH their mobility is already significantly reduced. Over the course of the presented here measurements, we decided to perform AFM scanning in the absence of salt-enriched buffer, which could lead to the crystallization of the salt on top of the membranes. Thus, membranes prepared in HEPES buffer, were thoroughly washed with MilliQ water to flush out remnants of salt. The absence of Na^+^ ions in the residual water leads to lower (<0.2 μm^2^/s) mobility of lipids under decreased hydration conditions, explaining the lack of macroscopic circularity changes within membranes at low humidity. However, it should be noted that even though in the dehydration conditions lipid mobility is affected, there was a noticeable structural reorganization in the partially dehydrated membrane compared to the membrane at fully hydrated conditions.

After dehydration, we performed gradual rehydration of the membrane to confirm the full preservation of the structure during the complete cycle of lowering and increasing the humidity. As presented in Fig. 3A and S4A throughout the entire rehydration process the integrity of lipid membranes remained undamaged. Iriarte-Alonso et al. reported that membranes composed of pure DOPC upon abrupt and complete dehydration and rehydration loose their structural arrangement and membrane continuity, resulting in abundance of various defects and holes, which exposed the bare support^31^. This gives even more importance to the dehydration method presented in our research, which allows the membrane and lipids to adapt to the slowly changing hydration state of the membrane. The high-resolution images of individual domains revealed increased mixing of the L_o_ and L_d_ phases as shown in Fig. 3B, and S4B. As shown in Fig. 3C the AFM height profiles across the domains showed that the height mismatch between the L_d_ and L_o_ phases the height mismatch between the L_d_ and L_o_ phase, was different for varying hydration conditions. Thus, we concluded that the more prominent mixing of phases at higher hydration values, resulted from two phenomena; (i) the gradual recovery of lipids mobility, leading to more dynamic membrane reorganization and (ii) increase of the hydrophobic mismatch between phases, which promotes the phase separation. Consequently, the L_d_ phase lipids separate from the _Lo_ phase and merge, to minimize the perimeter exposed to water (see Fig. S5).

Based on the AFM images we determined the overall bilayer thickness for the membrane without bulk water (90% RH) by selecting areas with membrane defects as shown in Fig. 4A. The formation of holes was induced by applying an abrupt dehydration procedure, which is based on the rapid aspiration of water with a pipette. Moreover, to find ruptured parts of the membrane, we have chosen areas near the edges of the solid substrate, which are more prone to air-water interfacial peeling force acting during dehydration. We extracted cross section profiles across the areas containing three types of features; holes, L_o_ and L_d_ phases and observed that the height difference between the L_d_ phase and the surface of bare substrate was 3.87 ± 0.21 nm, which we ascribed to the DMoPC bilayer thickness. It should be noted that this value is in agreement with the bilayer thickness of 3.86 nm reported by Lee et al. for a fully hydrated membrane also composed of DMoPC lipids_51_. From the height profiles, it was possible to distinguish domains of the L_o_ phase protruding approximately 1.3 nm from the Ld phase. Furthermore, the height profiles along the domains confirmed that the intermediate height areas represented the L_d_ phase trapped within the L_o_ domains, as the obtained height difference corresponds to the L_o_/L_d_ height mismatch (see Fig. 4B).

**Figure 4.**
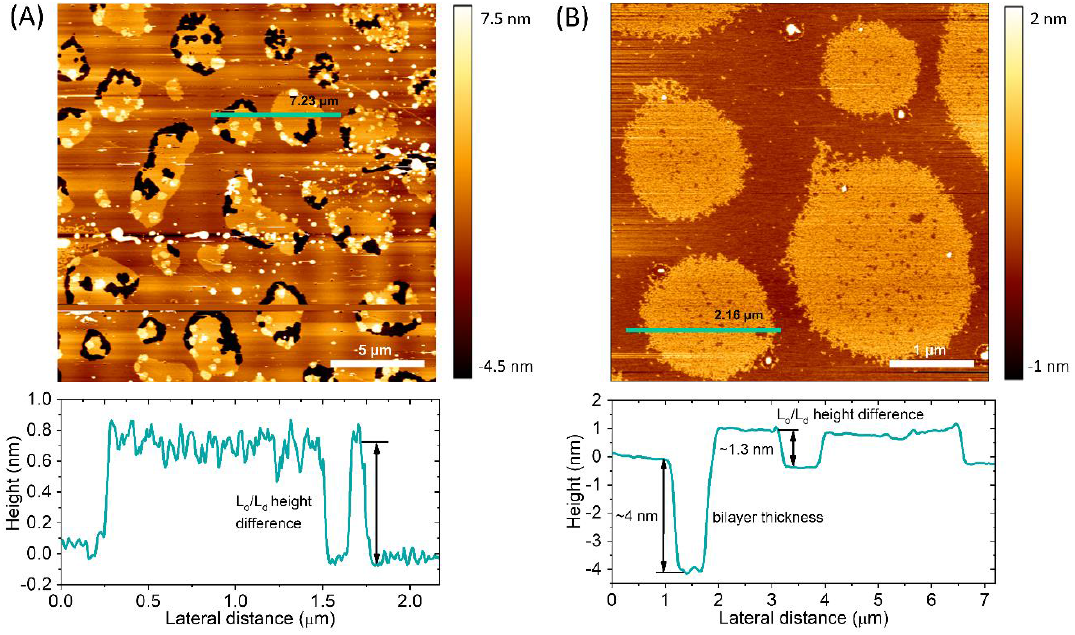
(A) SLB after abrupt removal of bulk water and consecutive equilibration at 90% RH. The bilayer thickness at 90% RH was around 4 nm with the height difference between the Lo and Ld phase was approximately 1.3 nm. (B) SLB after careful removal of bulk water and consecutive equilibration at 5% RH. The intermediate height areas within the Lo phase domains correspond to the Ld phase. The height difference between phases at this hydration conditions was approximately 0.8 nm.

### 3.3 Hydrophobic mismatch and line tension under dehydration conditions

Preparation of lipid membranes from the ternary lipid mixture used in this research leads to the formation of lipid domains composed of the L_o_ phase embedded within the Ld lipid matrix. AFM^52,53^ and X-ray scattering^54,55^ studies clearly show that the L_o_ phase, due to the presence of saturated and more densely packed lipids, is thicker than the L_d_ phase which contains unsaturated lipids. This leads to the so-called “height mismatch“ or “hydrophobic mismatch“ between the two phases. Although this height difference is well defined for SLBs composed of phospholipids with different fatty acid chain lengths in bulk water, it has not been measured in decreased hydration conditions due to the problems with maintaining membrane structural integrity upon desiccation^16^. From the height distribution histograms (see Fig. S6) we determined the height mismatch between phases in a broad range of membrane hydration states as shown in Fig. 5A. We observed that the height mismatch decreases from 1.36 ± 0.19 nm for 90% RH to 0.8 ± 0.04 nm for 5% RH during dehydration. These changes were almost fully reversible with rehydration. Importantly, the measurements in bulk water as well as in all hydration conditions were done with the same cantilever and in one mode (contact mode) to ensure that the obtained values are comparable.

**Figure 5.**
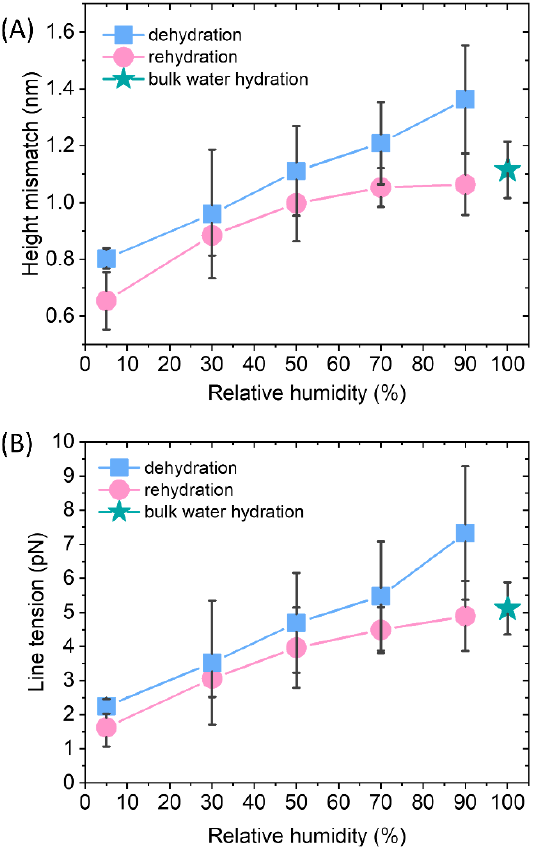
(A) Height mismatch between the L_o_ and L_d_ phases during dehydration and rehydration analysed from the height histograms of 3-8 different images, containing multiple domains of the Lo phase. (B) Line tension during dehydration and rehydration calculated from the model assuming presence of soft domains, no spontaneous curvature, and no changes in the height of the Lo phase during the hydration change cycle.

The lateral organization of the membrane, distribution of domains, their size and shape are all driven, among other, by the height mismatch between the L_d_ and L_o_ phase, which is a key factor leading to the occurrence of line tension at the boundary of two phases. The exposure of the lipids hydrophobic tails to the aqueous environment is energetically unfavorable. Thus, lipids of the L_o_ phase tend to organize themselves into domains to reduce the boundary at the interface of the phase, in consequence reducing the exposure to the aqueous medium. Theoretical model developed by Cohen et al. showed that the line tension increases quadratically with the phase height mismatch^14^. Experimental work based on this model has shown that by varying PC acyl chain length from 22 to 14 carbons, the height difference between phases increases from 0.17 ± 0.9 nm to 1.56 ± 0.13 nm, which leads to the 100-fold increase in the values of line tension, from 0.06 to 6 pN respectively^16^. Line tension with the similar order of magnitude has been obtained using also different approaches such as the measurement of domains nucleation rate^56^ or the analysis of vesicles geometry based on the Jülicher and Lipowsky theory^57^. Surprisingly, the line tension in membranes has never been calculated or measured as a function of its main determinant, which is the presence of water. From the extracted profiles for samples at 90% RH it could be concluded that the overall lipid bilayer thickness does not appear to change upon removal of bulk water, as the measured at 90% RH bilayer thickness 3.87 nm is consistent with the value 3.86 nm reported in the literature for fully hydrated DMoPC bilayer^51^. The determination of the L_o_ and L_d_ effective thickness was not possible for each hydration level, because lipid membranes did not exhibit any defects in the form of holes and membrane disruptions that would provide the reference plane for height measurements. While the induction of holes in the membrane is relatively easy as it requires abrupt removal of bulk water, once the membrane defects are formed and the hydration is further gradually decreased, they behave in an unpredictable manner; they rapidly extend their size in all directions, and the edges of the membrane curl up. This progressive degradation of membrane makes it hard to follow the absolute thickness of the bilayer during the changing of hydration. However, NMR and X-Ray diffraction experiments on membranes under lower hydration conditions showed that lipids forming the L_d_ phase are more prone to height changes upon dehydration. The L_d_ phase lipids undergo the straightening of acyl chains, which leads to a decrease in the lateral area per lipid molecule and consequently increase in their hydrophobic thickness^58,59^. Moreover, the recent simulations findings on the behavior of lipid membranes under decreased hydration clearly show that, in the absence of water, lipids from the L_d_ phase undergo liquid-to-gel crossover and exhibit characteristics of ordered membrane, which is more rigid with more densely packed lipids’ acyl chains^59^. Having this in mind, we infer that the change in height mismatch and consequent change in the line tension is caused by the straightening of the fatty acid chains of lipids composing the L_d_ phase, rather than by a modification in the thickness of the L_o_ phase. We calculated the line tension according to the equation proposed by Cohen et al.^14^ (see Materials and Methods section), assuming that the height difference was caused by the increase in the thickness of the L_d_ phase. We observed that there was a linear dependence of the line tension on the hydration level (Fig. 6B).

While all the above discussed evidence suggest that it is the L_d_ phase changing its height when controlling membrane hydration, we also considered two other scenarios, in which a decrease of the hydrophobic mismatch is caused by the reduction of the L_o_ phase thickness or by both phases undergoing the opposite effect, meaning the thickness of the L_d_ phase increases with the simultaneous decrease of the L_o_ phase thickness (see Fig. S7A). Regardless of the assumptions, for all three scenarios we observe the same trend in changes of the line tension under varying hydration level, that is an almost 3-fold decrease of the line tension for the lowest humidity of 5% RH when compared to 90% RH (see Fig. S7B and S7C).

## 4 Conclusions

In this research, we have presented a methodology for AFM measurements at varying hydration conditions in which biomimetic membranes are directly subjected to the slow, gradual changes of their hydration state. The phase-separated lipid membranes, characterized by the presence of the L_d_ and L_o_ phases, were exposed to a wide range of environmental humidity and at the same time, fluorescence images as well as information about membrane topography were acquired. We observed that the overall lateral organization of lipid membranes, that is the presence of phase separation and continuity of the structure, did not change upon removal of bulk water and subsequent, gradual dehydration of the membrane. However, the lack of bulk water leads to the increased miscibility of the lipids composing the L_d_ phase inside domains of the L_o_ phase, as compared to fully hydrated conditions. We addressed the response of biomembranes to the dehydration conditions in terms of the line tension, which was so far measured only as a function of lipids chain length and never through the direct impact of the water content. We observed a 3-fold difference in the line tension between the two extreme hydration states. The significant drop of the line tension at the boundary of the L_d_ and L_o_ phases explains the tendency of lipids from different phases to mix more freely under decreased membrane hydration.

Local dehydration of two merging cell membranes is an indispensable prerequisite in all fusion events, occurring during biological processes such as viral entry, endo- and exocytosis, neurotransmission or fertilization. Fusion can be modulated in two ways; through various types of fusogenic proteins, such as SNARE, or through the structural, local adaptation of the membrane lipid composition^60^. It has been shown that cholesterol accumulates at high curvature regions during stalk formation in fusion events^61^. This is ascribed to the ability of cholesterol to increase the membrane fluidity and induce formation of negative curvature. Sphingomyelin is known to promote formation of densely packed and less fluid L_o_ phase, effectively obstructing the membrane fusion^62^. The opposite effect is induced by unsaturated lipids forming L_d_ phase, which stimulate the membrane bending and promote formation of fusion intermediates^63^. Although, the merging of two membranes occurs preferably in high fluidity regions, viral protein elements such as HIV fusion peptides have been shown to bind to high line tension L_o_-L_d_ boundary regions, where they promote the membrane fusion of HIV viral envelope with the cell membrane of the host^64^. The revised stalk-pore model extended to heterogenous membranes (exhibiting L_o_-L_d_ phase coexistence) proposes that the reduction of line tension at the lipid phase boundary during stalk formation generates additional energy for the fusion, facilitating viral entry through phase boundary regions^65^. As we showed here, the lowering of the line tension under dehydration conditions is associated with an extensive migration of lipids between the L_d_ and L_o_ phase. It is thus feasible that the observed changes in local membrane fluidity and flexibility as well as the decreased line tension are yet another factor required for the fusion to occur. Altogether, our results provide new insights into the behavior of biological membranes under water scarcity conditions in terms of their lateral organization and adaptation to membrane dehydration. The findings presented here bring a new perspective to the processes that require local membrane dehydration such as cell fusion, fertilization, or binding of macromolecules. Finally, the proposed in this research methodology for AFM measurements under varying hydration states opens new possibilities for studying other biological systems and their interactions with water.

## Author Contributions

Emilia Krok: Conceptualization, Validation, Investigation, Writing -Original Draft, Visualization, Funding acquisition. Henri G. Franquelim: Conceptualization, Validation, Investigation, Resources, Writing -Review & Editing. Madhurima Chattopadhyay: Conceptualization, Validation, Writing -Review & Editing. Hanna Orlikowska-Rzeznik: Validation, Writing -Review & Editing. Petra Schwille: Writing -Review & Editing, Resources, Supervision. Lukasz Piatkowski: Conceptualization, Validation, Supervision, Writing -Original Draft, Funding acquisition.

## Acknowledgements

The authors acknowledge the financial support from the EMBO Installation Grant 2019 (IG 4147). LP acknowledges the financial support from the First TEAM grant No. POIR.04.04.00-00-5D32/18-00, provided by the Foundation for Polish Science (FNP). EK acknowledges the financial support from the Ministry of Education and Science of Poland in the year 2022 (Project No. 0512/SBAD/6212).

