## Supplementary data for "Nanoscale structural response of biomimetic cell membranes to controlled dehydration"

**Table of contents:**

Supplementary figures S1-S7

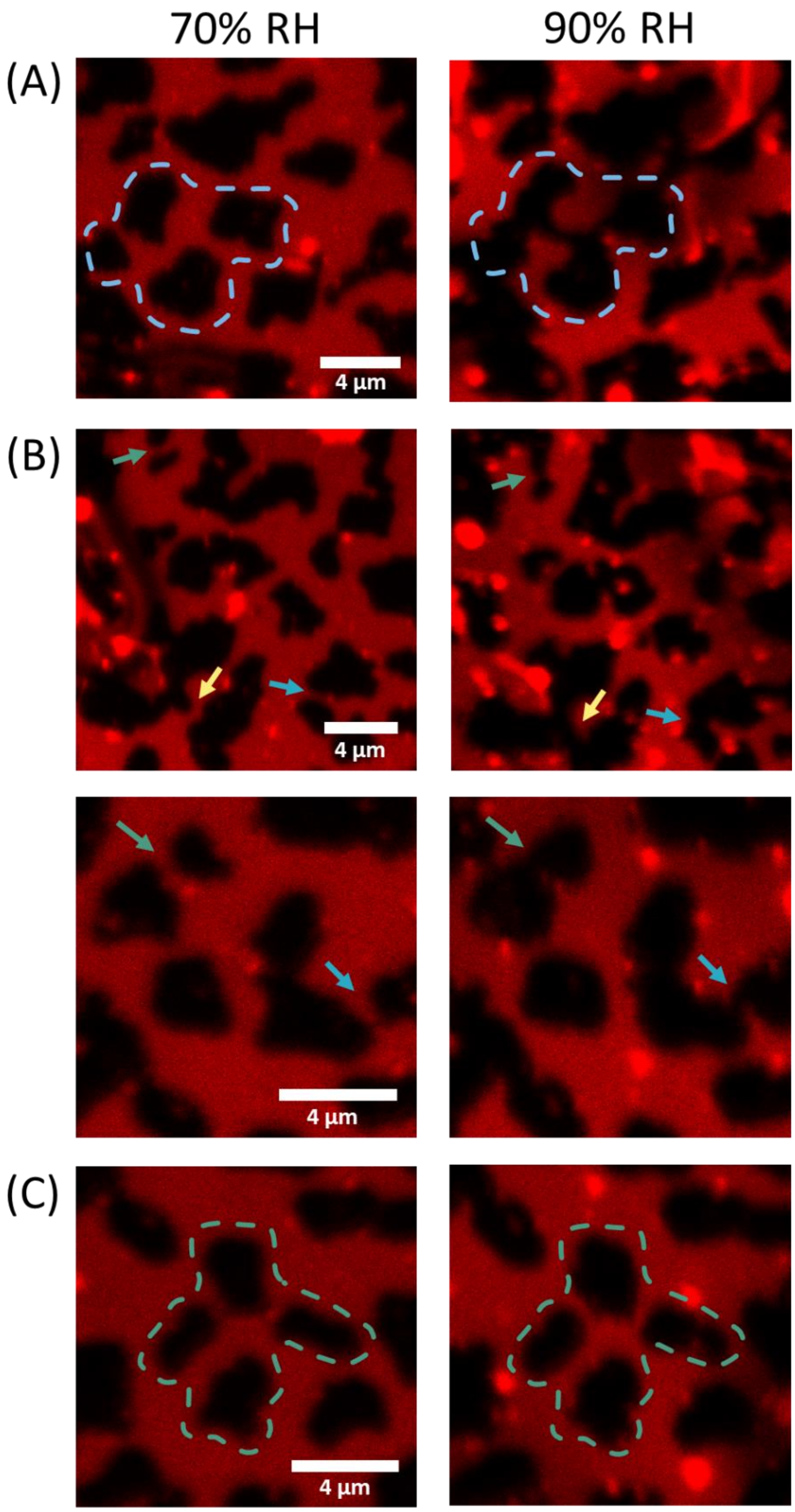


**Figure S1 Changes of the domains shape during rehydration**
(A) Merging of domains when the relative humidity is increased from 70 to 90% RH. The blue dashed lines mark the outlines of four merging domains. (B) Merging of domains during increase of humidity, arrows point out the merging points of the domains at 70 (left) and 90% RH (right). (C) Domains that did not merge during increase of the environment relative humidity. Confocal imaging was performed in 10 mM HEPES and 150 mM NaCl buffer.

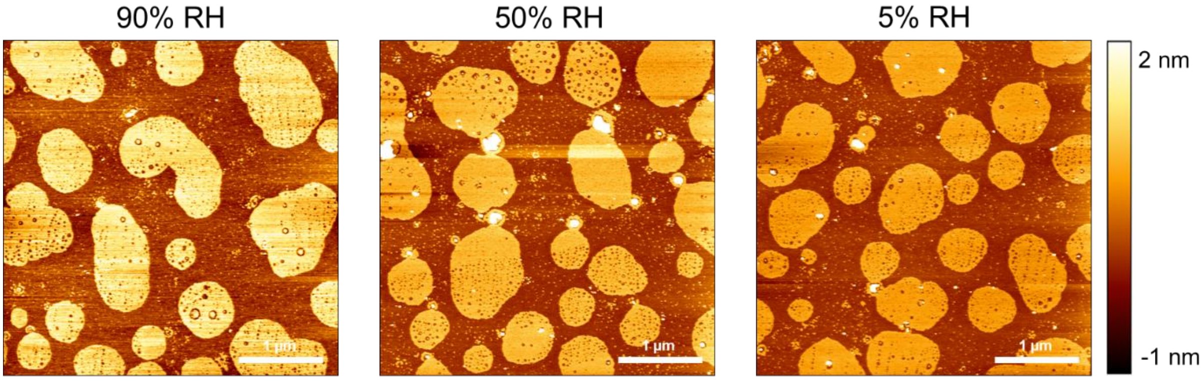


**Figure S2 AFM topography images of SLBs during dehydration**
Representative images of SLBs during dehydration cycle. Membrane was equilibrated to 90, 50 and 5% RH.

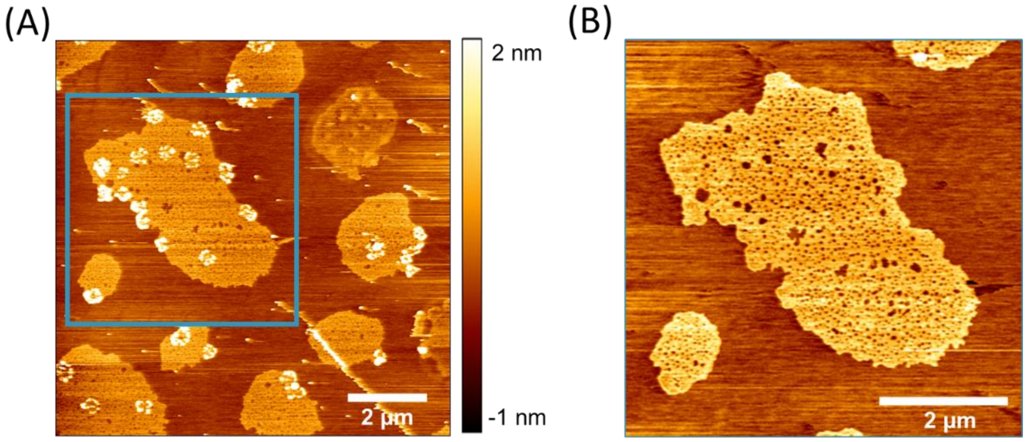


**Figure S3 AFM topography images of SLBs right after removal of bulk water**
(A) Representative image of lipid bilayer right after removal of bulk water and equilibration to 90% RH. Blue square indicates the area presented in high resolution images of lipid bilayer at 90% RH (B). Lipid membrane at high humidity was sticky, small aggregates on top were dragged by the tip, leading to distortions in the measurements. (B) After further equilibration and sweeping with the AFM tip over the surface, the aggregates on top of the membrane were removed and the topography could be measured more reliably.

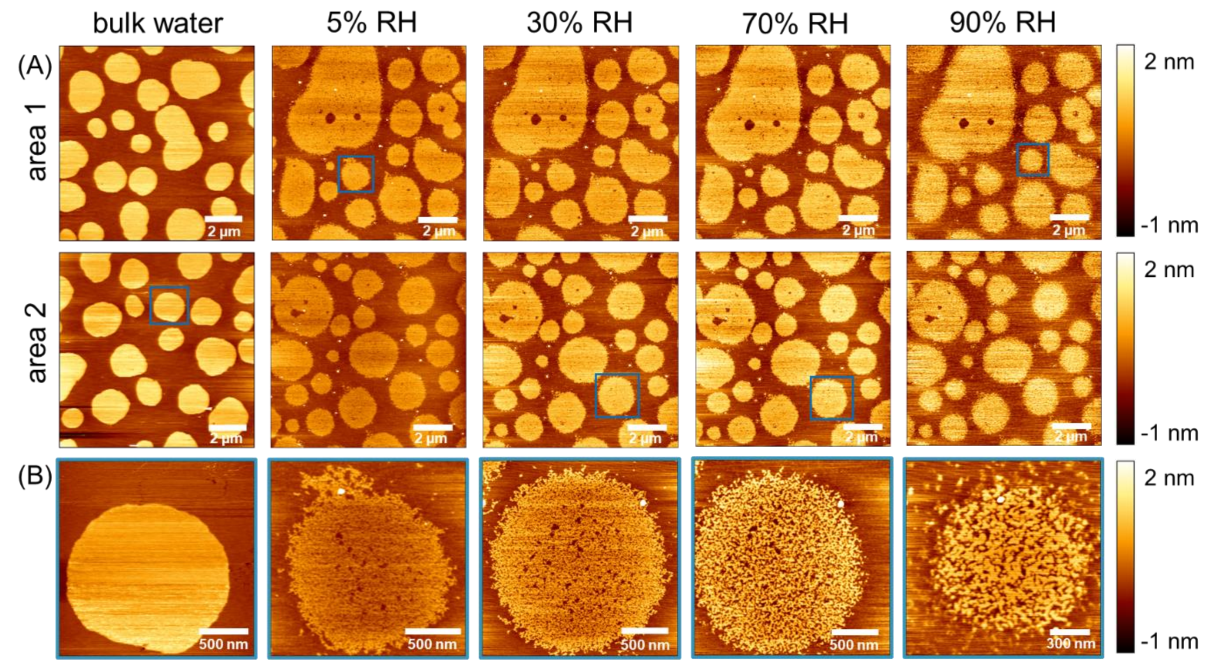


**Figure S4 AFM topography images of SLBs at different hydration levels.**
(A) Representative images of fully hydrated SLB, and SLB after removal of bulk water and equilibration to 5, 30, 70 and 90% RH. The top and middle rows correspond to two different areas of the SLB. (B) High resolution images of individual domains (indicated in panel A with blue squares) at different hydration levels.

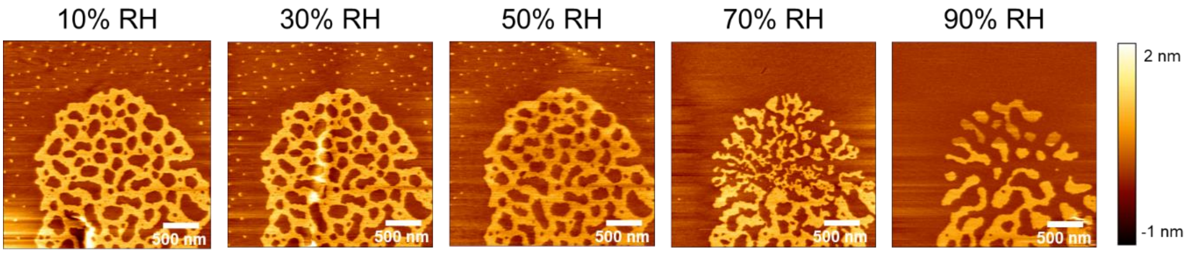


**Figure S5 High resolution AFM topography images of single $L_o$ phase domain at different hydration levels.**
High resolution images of single domain during membrane rehydration revealed evolution of the $L_d$ nanodomains trapped within the $L_o$ phase. At 70% RH it is evident that $L_d$ phase nanodomains merge to minimize the boundary between phases.

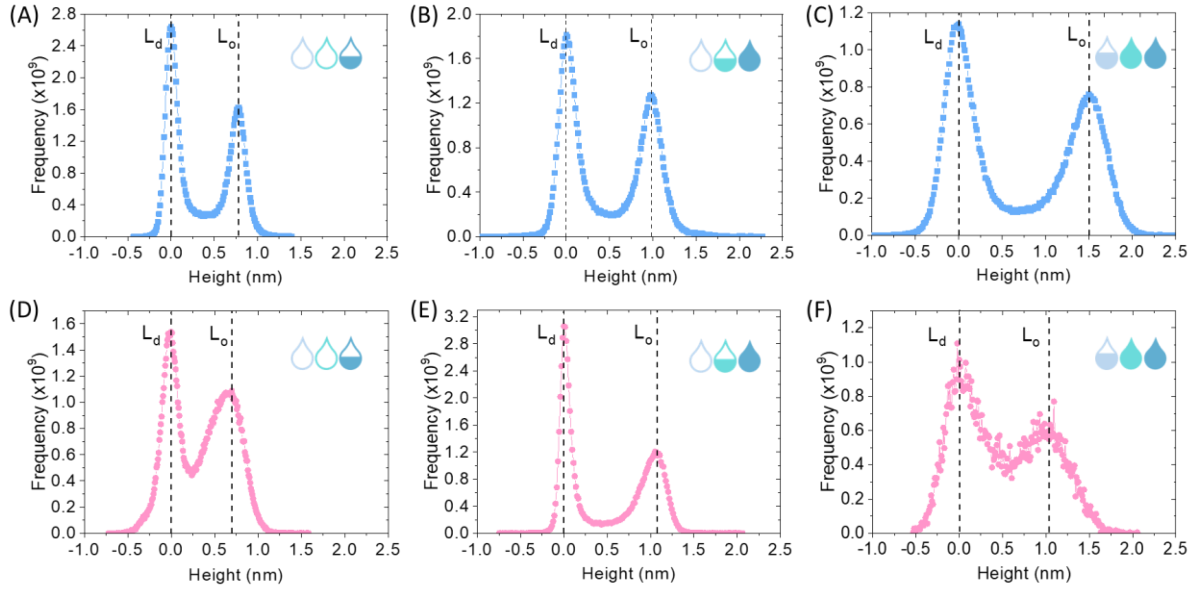


**Figure S6 Quantification of the height difference between the $L_d$ and $L_o$ phase from the AFM images.** Height distribution histograms for the membrane subjected to dehydration (top row, blue) at (A) 5% RH, (B) 50% RH, and (C) 90% RH, and rehydration (bottoms row, pink) at (D) 5% RH, (E) 50% RH, and (F) 90% RH. Profiles were offset corrected resulting in the $L_d$ peak being centered at 0 nm.

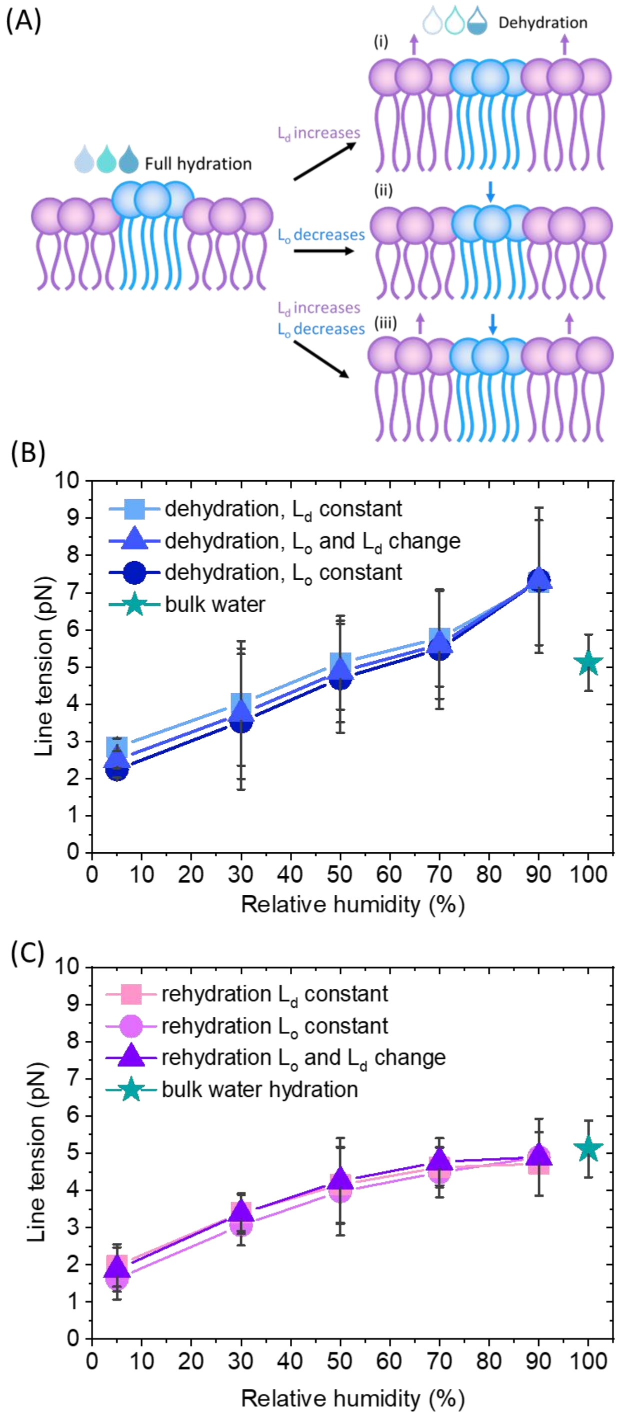


**Figure S7 Line tension at different hydration levels during dehydration and rehydration of the lipid bilayer.** (A) Three scenarios for the line tension calculation: (i) the $L_d$ phase thickness increases, while the $L_o$ phase does not change, (ii) the $L_o$ phase thickness decreases, while the $L_d$ phase thickness does not change, (iii) the $L_d$ phase increases and the $L_o$ phase decreases. (B) Line tension during dehydration calculated from the model assuming a soft domain and no spontaneous curvature for the three different scenarios presented in (A). (C) Line tension during rehydration calculated for the three different scenarios presented in (A).
